# Interaction between native and prosthetic visual responses in optogenetic visual restoration

**DOI:** 10.1101/2024.12.28.630611

**Authors:** Eleonora Carpentiero, Steven Hughes, Jessica Rodgers, Nermina Xhaferri, Sumit Biswas, Michael J Gilhooley, Mark W Hankins, Moritz Lindner

## Abstract

Degenerative retinal disorders leading to irreversible photoreceptor death are a common cause of blindness. Optogenetic gene therapy aims to restore vision in affected individuals by introducing light sensitive opsins into the surviving neurons of inner retina. While up until now the main focus of optogenetic therapy has been on terminally blind individuals, treating at stages where residual native vision is present could have several advantages. Yet, it is still unknown how residual native and optogenetic vision would interact if present at the same time.

Using transgenic mice expressing the optogenetic tool ReaChR in ON-bipolar cells, we herein examine this interaction through electroretinography (ERG) and visually evoked potentials (VEP). We find that optogenetic responses show a peculiar ERG signature and are enhanced in retinas without photoreceptor loss. Conversely, native responses are dampened in the presence of ReaChR. Moreover, in VEP recordings we find that optogenetic responses reach the cortex asynchronous to the native response.

These findings should be taken into consideration when planning future clinical trials and may direct future preclinical research to optimize optogenetic approaches for visual restoration. The identified ERG signatures moreover may serve to track treatment efficiency in clinical trials.

## INTRODUCTION

Degenerative retinal disorders are among the commonest causes of vision loss in industrial countries and represent a major socioeconomic burden [1–4]. These include more widespread conditions such as multi-factorial Age-related Macular Degeneration (AMD) and the rare ones such as Inherited Retinal Degenerations (IRDs, e.g. retinitis pigmentosa). Until now, more than 320 genes for retinal diseases have been identified [5]. While pathogenically diverse, these degenerative retinal disorders commonly lead to irreversible loss of rod and cone photoreceptors. Pharmacological and gene therapy-based approaches to slow down or halt disease progression are currently emerging [6, 7], yet each of these are only applicable to one particular condition. Moreover, they are not effective once photoreceptors have been lost [8].

It has long been proposed that ectopic expression of light sensitive proteins in inner retinal neurons, which survive during retinal degeneration, could render these neurones directly light sensitive and thereby substitute the function of the lost photoreceptors. Using this strategy, which is termed *optogenetic visual restoration*, light perception has been restored in animal models as well as in a patient with end-stage retinitis pigmentosa [9–16].

Currently, most optogenetic vision restorative approaches are performed in end-stage retinal degeneration. Yet, there are several reasons why an earlier intervention might be preferable. First of all, while primarily affecting the outer retina, retinal degeneration is accompanied by remodelling and rewiring of inner retinal circuity [17]. This rewiring might hinder orthodox intraretinal signal processing, which is a requirement for complex image forming vision and thereby limit the functional outcome of optogenetic vision restorative therapies. Therefore, it is conceivable that early optogenetic intervention could not only provide better functional results soon after treatment, it might also prevent further retinal remodelling and thereby deliver better long-term outcome. Secondly, offering optogenetic therapy only to those with end-stage retinal degeneration would mean leaving individuals affected by retinal degeneration with low vision, insufficient for daily routine tasks, for over years before they finally become eligible for an optogenetic therapy. Third, in macula disease, “end-stage retinal degeneration”, i.e. a situation without residual native light perception, is never reached. Macular diseases, the commonest being AMD, however, far outnumber the cases of rod-cone degenerations [3, 4] and restricting the availably of optogenetic gene therapy to terminally blind individuals would exclude this large collective of patients.

Especially in AMD it becomes clear that disease progression is a trajectory in time and in space, with outer retinal atrophy starting at one location on the macula and then spreading over time [18, 19]. Thus, areas with photoreceptor loss and therefore no native light perception would neighbour areas of intact retina with functional photoreceptors (**Figure 1**). While only the first would be the target of an optogenetic therapy, with current methods of gene delivery, optogenetic tool expression also in the neighbouring retina would be unavoidable.

**Figure 1:**
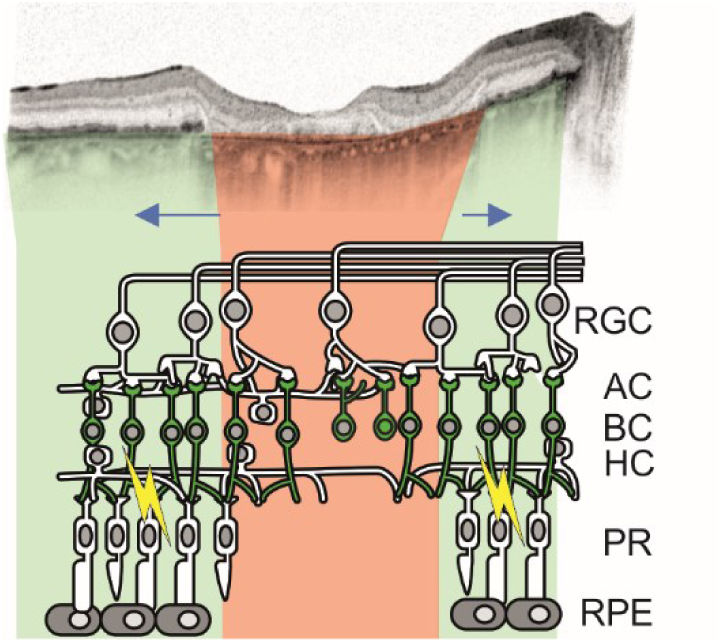
Optogenetic vision restoration in a retina with residual native visual function, exemplified on a case of atrophic age-related macular degeneration. Top: Optical coherence tomography of the macula of an eye of an 81-year-old female patient showing Geographic Atrophy secondary to Age-related macular degeneration [modified from: 8]. Highlighted in red are areas where retinal pigment epithelium (RPE) and photoreceptor (PR) atrophy is present. Inner retinal remodelling would occur inside these areas over time. In green, areas with preserved RPE and PRs. Bipolar cells (BC), likely preferred target of an optogenetic gene therapy, are highlighted in green. Optogenetic vision and residual native vision could possibly interfere in areas with intact PRs (indicated by yellow lightnings). Blue arrows indicate spatial progression of retinal degeneration over time.

Thus, optogenetic intervention early in the disease course would mean that residual native and optogenetic vision co-exist and, in some way, interact. The consequences for visual processing and perception are yet unknown. It is therefore necessary to understand this interaction in order to predict the impact of optogenetic tool expression in remaining/functional areas of the retina, and thereby evaluate how early intervention trials are safe in humans from a functional perspective.

In the present paper, we perform electroretinogram (ERG) and visually evoked potentials (VEP) recordings from transgenic mice expressing the optogenetic tool ReaChR in ON-bipolar cells (OBC) in the presence of native photoreceptors. We find that optogenetic responses show a peculiar ERG signature and are enhanced in retinas without photoreceptor loss. Conversely, native responses are dampened in the presence of ReaChR. We also find that optogenetic responses reach the cortex in advance to the native response potentially complicating perception. These findings should be taken into consideration when planning future clinical trials and may direct future preclinical research to optimize optogenetic approaches for visual restoration. The identified ERG signatures moreover may serve to track treatment efficiency in clinical trials.

## RESULTS

### A transgenic model to study the interference of native and prosthetic vision

We have previously described a mouse model expressing the long-wavelength-activatable variant of channelrhodopsin 2, ReaChR [20] in ON bipolar cells of retina-degenerate mice carrying the rd1 mutation in the Pde6b gene [11]. In these mice ON bipolar cell specific ReaChR expression is achieved by Cre-recombinase expression under the control of OBC specific Grm6 promoter [21]. In order to create a mouse model that would have both, native and optogenetic/prosthetic light responses, we bred out the Pde6b^rd1^ alle by crossing against C57Bl/6J mice. We thereby obtained three different genotypes of mice that are used throughout this study: 1) natively sighted, Pde6b^wt/wt^.Grm6^wt/wt^.*ReaChR* mice that do not express ReaChR (“wild-type” [“wt”] hereafter), 2) natively and optogenetically “sighted”, Pde6b^wt/wt^.Grm6^iCre/wt^.*ReaChR* mice that express ReaChR in OBC (“wt+ReaChR” hereafter) and 3) natively blind and optogenetically “sighted”, Pde6b^rd1/rd1^.Grm6^iCre/wt^.*ReaChR* mice that express ReaChR in OBC (“rd1+ReaChR” hereafter). Median age of mice used in this study was 75 [IQR: 62 - 79] days for wt, 75 [IQR: 74 - 81] days for wt+ReaChR and 75 [IQR: 71 - 78] days for rd1+ReaChR. We confirmed the expected expression patterns as well as the presence and absence of native photoreceptors using immunohistochemistry (**Figure 2**).

**Figure 2:**
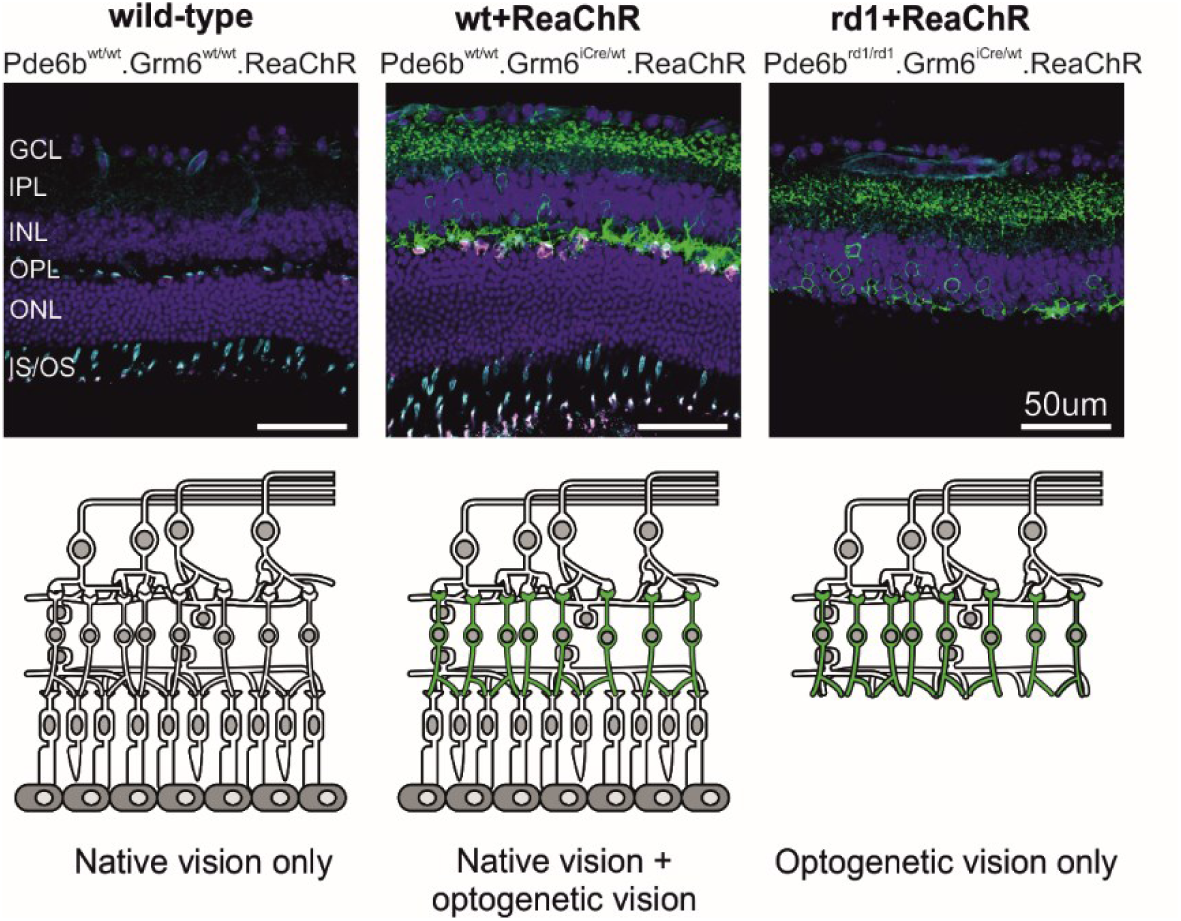
Schematic overview and immunohistochemistry for the retina of the mouse lines used in this study. Top row: representative confocal micrographs of retinal cryosections from a “wild type” (Pde6b^wt/wt^.Grm6^wt/wt^.ReaChR) mouse (left), a ReaChR-expressing non-degenerate (Pde6b^wt/wt^.Grm6^iCre/wt^.*ReaChR*) mouse (middle), and ReaChR-expressing retina-degenerate mouse (Pde6b^rd1/rd1^.Grm6^iCre/wt^.*ReaChR*, right). Sections were co-lablled with anti-GFP (green), PNA Lectin (cyan), anti-Cone-Arrestin (magenta), and DAPI (blue). Bottom row: Corresponding schematic representations. The green colour illustrates ReaChR-expression in ON-bipolar cells. IS/OS: Inner Segment/Outer Segment, ONL: Outer Nuclear Layer, OPL: Outer Plexiform Layer, INL: Inner Nuclear Layer, IPL: Inner Plexiform Layer, and GCL: Ganglion Cell Layer

### Interaction of native and optogenetic light responses

To assess if the expression of an optogenetic tool in the retina would interact with native visual responses we commenced our functional study with ERG recordings from wt+ReaChR mice and wt mice not expressing ReaChR. We first recorded scotopic and mesopic responses from dark-adapted mice to 0.01 cd×s/m^2^ and 3 cd×s/m^2^ flash stimuli, respectively. Both a- and b-wave responses observed in wt+ReaChR mice were statistically indifferent from those observed in wt mice. Yet, there was a minor trend towards longer b-wave implicit times for 3 cd×s m^2^ wt+ReaChR mice (wt: 35.75 [IQR: 35.5 – 37.88] ms, n=6, vs. wt+ReaChR 39.5 [IQR: 38.5 – 41.5] ms, n=7, p = 0.056; **Figure 3**, **Supplementary Table 1**). For comparison, 3 cd×s/m^2^ ganzfeld illumination would result in a photon flux of approximately 3 × 10^13^ photons/cm^2^/s on retinal level, which is substantially below the values that have been reported necessary to induce ReaChR mediated chances in neuronal activity [11].

**Figure 3:**
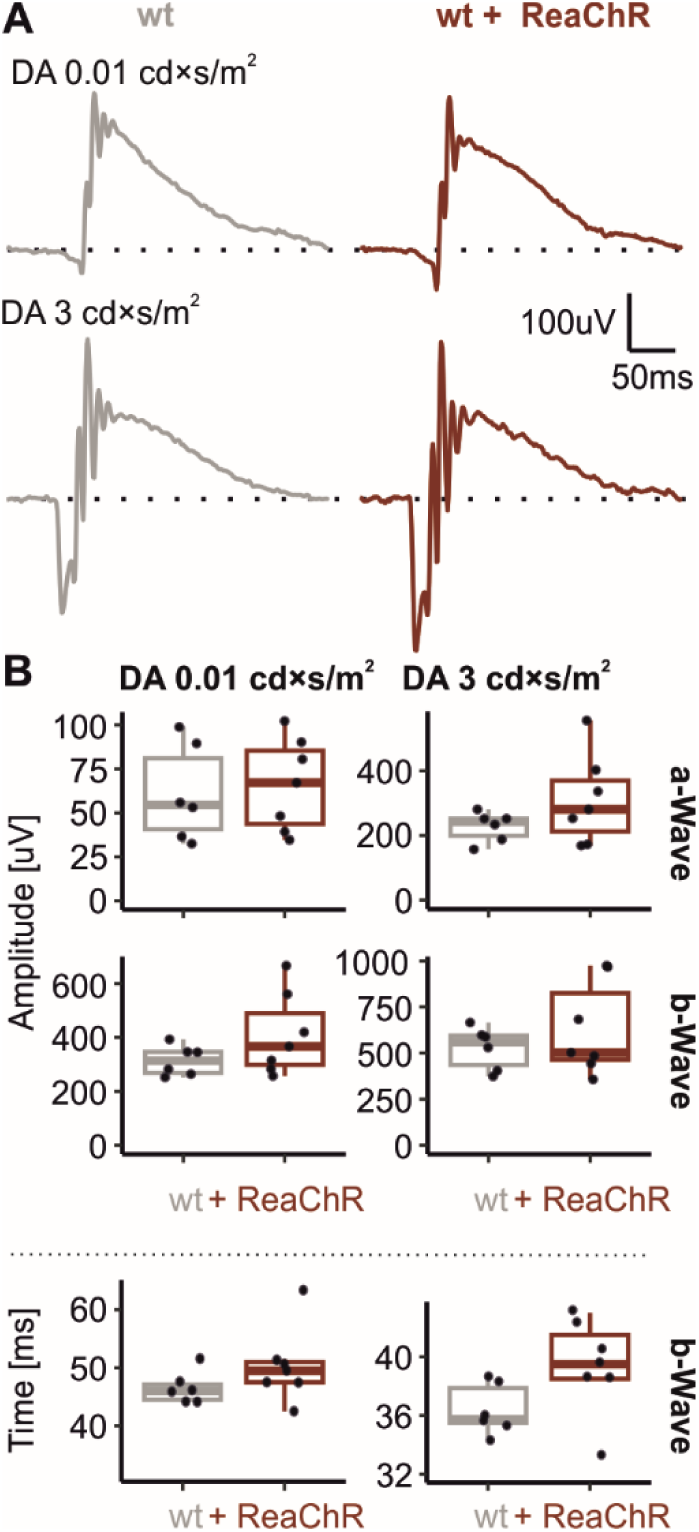
Dark-adapted (rod-dominated) ERG recordings are not altered in mice expressing ReaChR in ON-bipolar cells. (A) Representative recordings obtained in response to 0.01 cd × s / m^2^ and 3 cd × s / m^2^ 4 ms flash stimuli from wild-type mice (grey) and ReaChR-expressing, non-degenerate mice (dark red). (B) Summary statistics for a- and b-waves amplitudes and implicit times as obtained in response to those stimuli.

We continued by assessing photopic responses in light-adapted mice exposed to a series of flash stimuli of increasing energy. Therefore, mice were exposed to 30 cd/m^2^ for at least 8 minutes before the beginning of the recording and all subsequent stimuli were delivered on that background. ERG waveforms exhibited the normal shape expected for cone-mediated responses in mice up to a stimulus energy of 30 cd×s/m^2^ in both genotypes. For comparison, no light response could be observed in rd1+ReaChR mice for these stimuli (**Figure 4 A**). At flash stimulus energies of 100 cd×s/m^2^ and above, an early and rapid negative deflection appeared before the onset of the a-wave in wt+ReaChR mice, but not in wt mice (**Figure 4 A, red arrows**). From the same stimulus energy doses onwards, rd1+ReaChR showed a similarly early and fast deflection. A stimulus energy of 100 cd×s/m^2^ on ganzfeld illumination would result in a photon flux of approximately 1.17 × 10^15^ photons/cm^2^/s on retinal level, which matched the intensity threshold for ReaChR mediated changes in neural activity observed in earlier studies [11]. We therefore hypothesized that this early negative deflection is a consequence of ReaChR activation and therefore termed it a_o_-wave (with “o” for “optogenetic”). In line with this assumption, we observed an even larger a_o_ amplitude for 900 cd×s/m^2^ in wt+ReaChR mice, while the a-wave amplitude as observed in wt mice did not increase further, nor did any additional a_o_-like deflection appear (**Supplementary Figure 1**). In wt+ReaChR mice a_o_ peaked at 8.0 [IQR: 7.1 - 8.8] ms for a 100 cd×s/m^2^ stimulus, which was significantly faster than the implicit time of the a-wave observed in wt mice (17.4 [IQR: 16.3 - 21.3] ms, p < 0.001, (**Figure 4 B**). The onset of a_o_ occurred virtually instantaneously, with the first data point (0.5 ms, on the filtered ERG traces) after stimulus onset already showing a clear deviation from the zero line. Interestingly, comparing a_o_ amplitudes between wt+ReaChR and rd1+ReaChR mice, we observed that that a_o_ amplitudes were substantially larger in wt+ReaChR as compared to rd1+ReaChR (wt+ReaChR: 33.76 [IQR: 23.14 - 46.31] uV, n=11, vs. 6.36 [IQR: 5.13 - 7.66] uV, n=6, p < 0.001; **Figure 4 C**, **Supplementary Table 1**).

**Figure 4:**
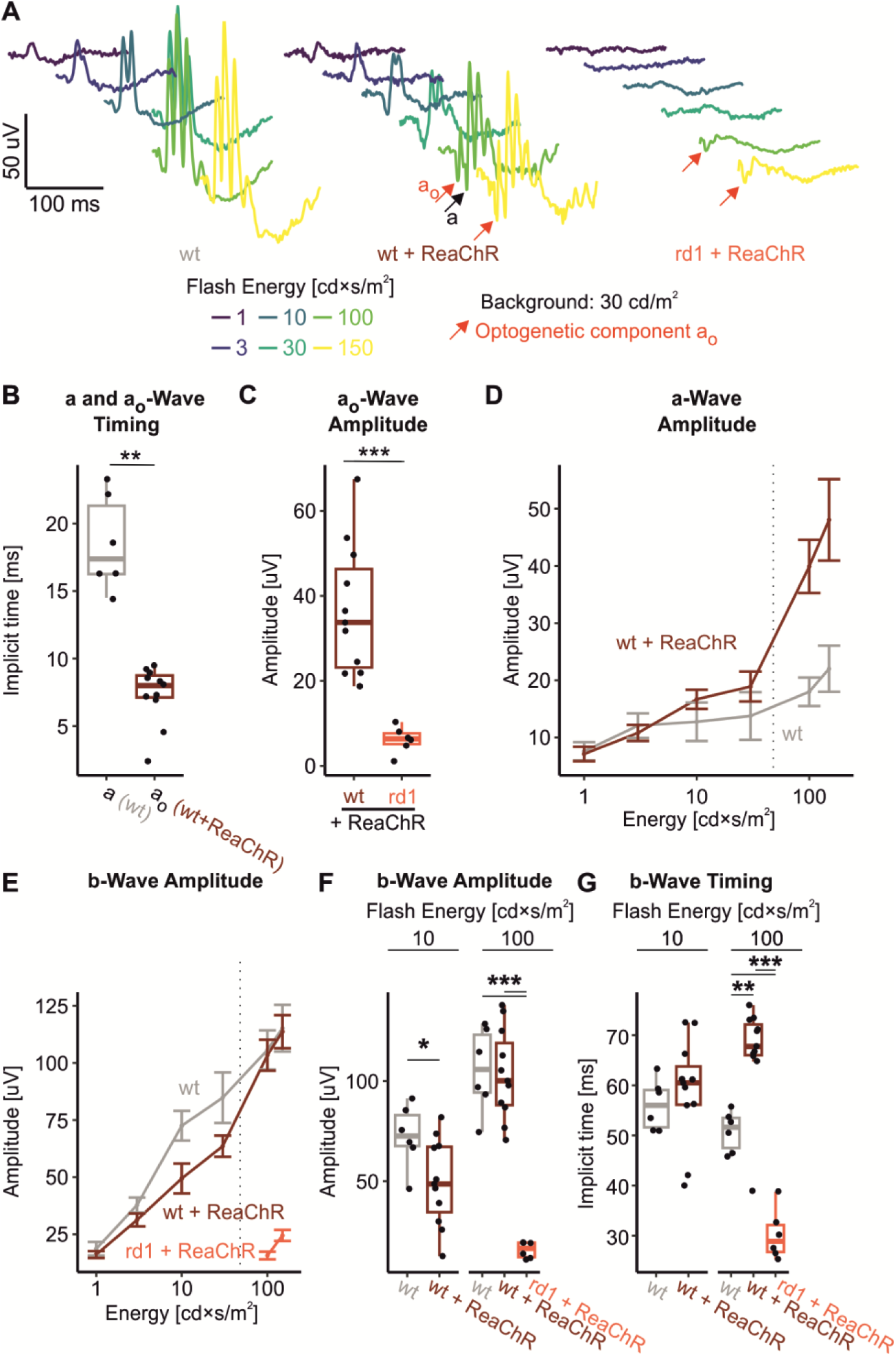
Light adapted ERG recordings. **(A)** Representative recordings for increasing flash stimulus energies from wild-type mice (left), ReaChR-expressing, non-degenerate mice (middle) and ReaChR-expressing, retina-degenerate mice (right). Note the appearance of a novel, putatively ReaChR-driven ERG component “a_o_”. **(B, C)** Box-plots comparing (B) the implicit times for a-waves (measured in wild-type mice) and a_o_-waves (measured in ReaChR-expressing, non-degenerate mice) and (C) a_o_ amplitudes in ReaChR-expressing, non-degenerate and ReaChR-expressing, retina-degenerate mice in response to a 100 cd × s / m^2^ flash. **(D, E)** Stimulus - response curves for a- and b-wave amplitudes, respectively. B-wave amplitudes presented in D are measured relative to the a-wave. Dotted line indicates the “activation threshold” for ReaChR (8 × 10^14^ photons/cm^2^/s = 68 cd × s / m^2^), i.e. the minimum stimulus energy required to induce a change in spike firing rate as determined in: [11]. As the effects on a-and b-wave amplitudes observed where opposing in direction we additionally calculate net b-wave amplitudes (i.e. relative to the zero line) for reference (**Supplementary Figure 2**, **Supplementary Table 1**).**(F)** Corresponding summary statistics for b-wave amplitudes at 10 cd × s / m^2^, where no major ReaChR activation would be expected, and 100 cd × s / m^2^, where both, native cone-opsins and ReaChR are expected to be activated. **(G)** b-wave implicit times for the same stimulus conditions as in (F).

We next focused on analysing the typical ERG components, in particular a- and b-waves. In case of a-wave amplitudes, we observed a minimal increase in amplitudes for 10 cd×s/m^2^ and 30 cd×s/m^2^ stimuli in wt+ReaChR mice as compared to wt mice, and this increase became more robust from 100 cd×s/m^2^ on (**Figure 4 D**). a- and a_o_-wave amplitudes did correlate (r = 0.767, p < 0.01, n = 9), and a_o_-waves had a rather broad trough in the rd1+ReaChR mice, suggesting that the steep increase measured for a-wave amplitudes from 100 cd×s/m^2^ on might at least partially reflect a superposition of the a-wave and the end of the preceding a_o_-wave. For the b-wave – in turn – we observed a dampening of response amplitudes for the 10 cd×s/m^2^ and 30 cd×s/m^2^ stimuli (for 10 cd×s/m^2^, wt: 61.92 [IQR: 51.29 - 69.02] uV, n=6, vs. wt+ReaChR: 34.41 [IQR: 23.4 - 46.94] uV, n=11, p<0.01; **Figure 4 E and F**, **Supplementary Table 1**).

With regard to timing, we saw the same (non-significant) tendency towards longer b-wave implicit times 10 cd×s/m^2^ in the wt+ReaChR group that we had already observed under dark-adapted conditions. For 100 cd×s/m^2^ stimuli this trend became substantially more pronounced, and the difference towards wt mice not expressing ReaChR became statistically significant (51.62 [IQR: 47.5 - 53.5] ms, n=6, vs. 67.75 [IQR: 66 - 72.12] ms n=11, p<0.001; **Figure 4 G**, **Supplementary Table 1).**

### Nature of the b-wave dampening

B-wave dampening could either reflect a true interaction of the optogenetic and native visual responses at cellular level or it could be a simple superposition of the field potentials caused by the parallel stimulation of ReaChR and the native opsins. The latter needs to be considered as ReaChR is expressed cone- and rod-OBC [11], whereof rod-OBC would not be directly activated by our cone directed stimuli. To address this possibility, we developed an ERG stimulus paradigm that would allow temporally independent stimulation of ReaChR and cone opsins. ReaChR does not inactivate (completely) and is maximally sensitive to light in the yellow/red spectrum (λ_max_=590 nm) [20]. We therefore chose to deliver blue (LED centre wavelength: 455 nm) light flashes of moderate intensity (5 cd×s/m^2^) on an amber (Red + Green primaries, CIE(x) =0.528, CIE(y) =0.427, 3500 cd/m^2^) background. This amber background would be bright enough to evoke weak continuous ReaChR activation, while cones – in particular S-cones, that have their spectral sensitivity maximum in the blue range – would adapt to this background lighting and then strongly respond to the incident flash of blue light. Comparing b-wave amplitudes between wt and wt+ReaChR mice that were obtained in response to such a stimulus paradigm, we observed a significant reduction of amplitudes in wt+ReaChR to 53.8 % of the wt values (wt: 107.86 [IQR: 77.25 - 140.37] uV, n = 8, vs. wt+ReaChr: 58.09 [IQR: 52.63 – 73.76] uV, n = 11, p< 0.05; **Figure 5 A and B**, **Supplementary Table 1**). Thereby, the b-wave dampening was more pronounced upon ReaChR-pre-activation than it was without such pre-activation (67.0 % for 10 cd×s/m^2^ on ISCEV-standard 30 cd/m^2^ background, see above, **Figure 4 E and F**), strongly indicating that b-wave dampening reflects a true biological interference of the optogenetic and native signals within the same cells.

**Figure 5:**
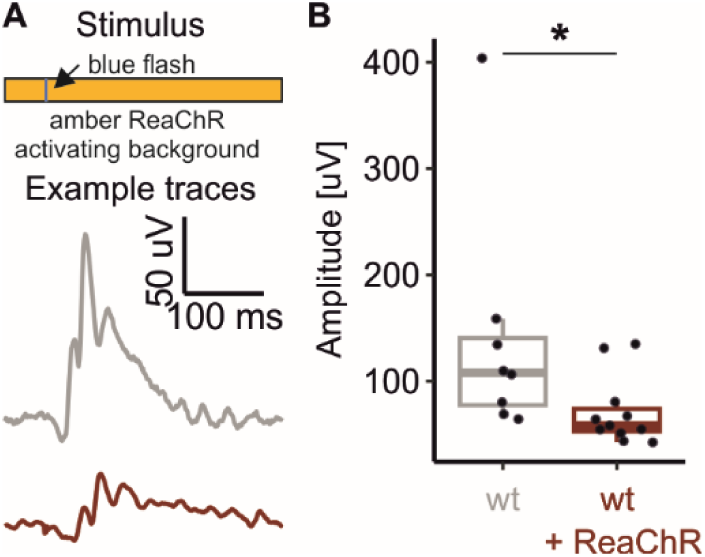
S-cone mediated light-responses are dampened on a ReaChR activating background. **(A)** Illustration of the stimulus protocol: 4 ms 5 cd × s / m^2^ blue flash stimulus on an amber 3500 cd/m^2^ background) and exemplary responses recorded from wild-type (grey) and ReaChR expressing, non-degenerate mice (dark red). **(B)** Summary statistics for the obtained b-wave amplitudes. Suppression of b-wave responses is ReaChR expressing mice is more pronounced than observed in response to a 10 cd × s / m^2^ flash stimulus on a 30 cd / m^2^ background as shown in Figure 3 C and D.

Thus, dampening of b-waves is presumably due to attenuation of the biophysical properties of OBCs, where optogenetic and native visual electrical responses converge. Yet, an alternative hypothesis is that the expression/activation of ReaChR in OBCs primarily affects inner retinal computation, e.g. by affecting the interplay between bipolar cells, horizontal cells and amacrine cells, which then leads to feedback on OBCs and finally b-wave dampening. To address this possibility we analysed the oscillatory potentials that reflect this inner retinal signal processing (predominantly inhibitory amacrine feedback) [22]. We did so for a stimulus energy where b-wave dampening but no a_o_-wave was observed (10 cd×s/m^2^) as well as for a stimulus energy strong enough to induce an a_o_-wave (100 cd×s/m^2^, **Figure 6 A and B**). Oscillatory potentials were indifferent in terms of the frequency of their principal spectral component (**Figure 6 C**) as well as the spectral power at that frequency (**Figure 6 D**, **Supplementary Table 1**). This suggests that the b-wave dampening seen in wt+ReaChR mice is rather due to a process upstream to inhibitory amacrine feedback, supporting the concept of an attenuation of the biophysical properties of OBCs.

**Figure 6:**
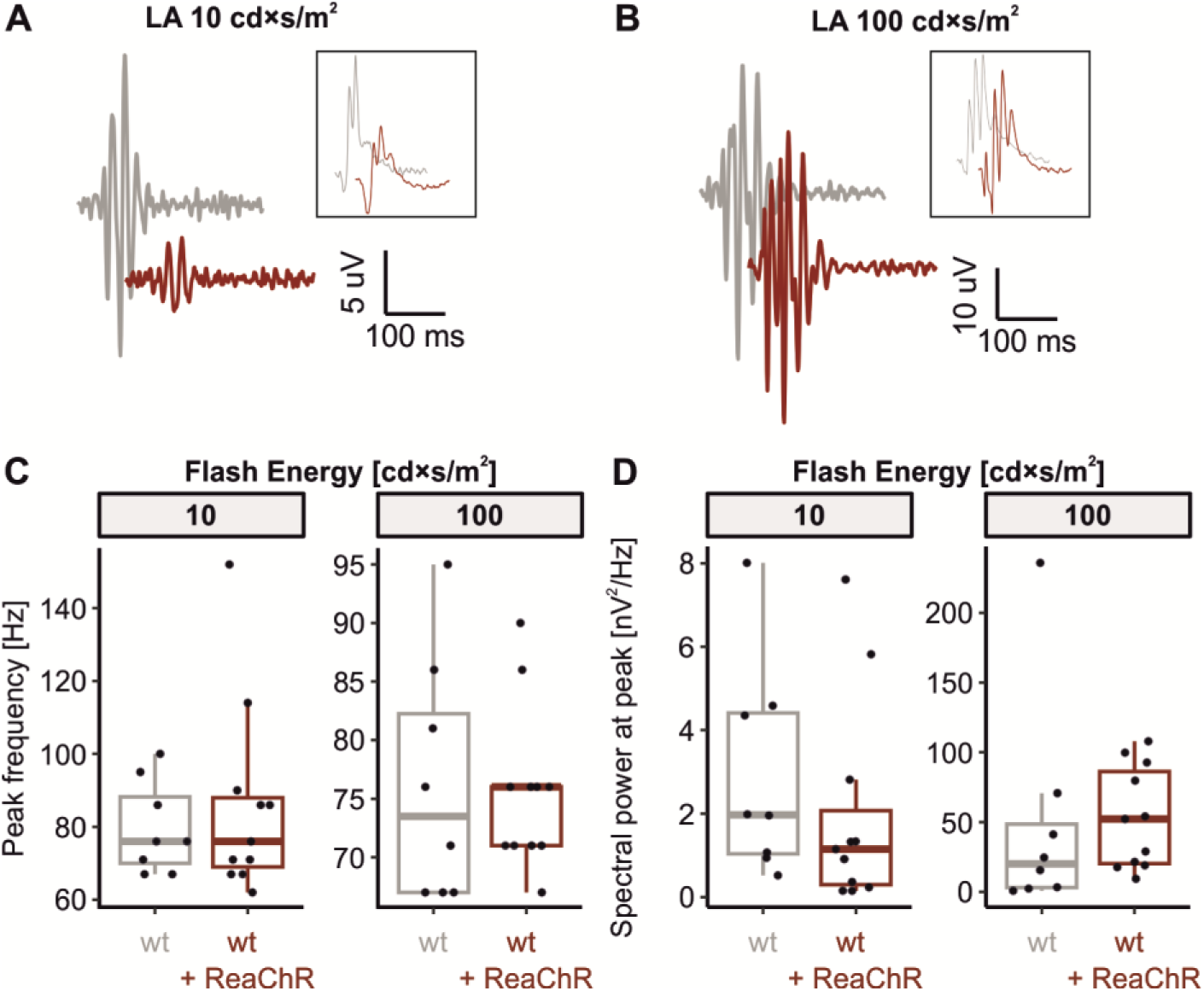
Photopic oscillatory potentials in light adapted ERG recordings are not altered in mice expressing ReaChR in ON-bipolar cells. **(A, B)** Representative recordings obtained in response to 10 cd × s / m^2^ and 100 cd × s / m^2^ flash stimuli from wild-type mice (grey), ReaChR-expressing, non-degenerate littermates (dark red). Shown are the oscillatory components (filter bandpass 75 – 300 Hz) as well as the originating ERGs (filter bandpass 0.5 - 300 Hz, insets, not in scale). **(C, D)** Summary statistics for the peak frequency of the oscillatory potentials **(C)** as well as the spectral power at that frequency **(D)**. Data are shown for 10 cd × s / m^2^ flash stimuli, where no major ReaChR activation would be expected, and 100 cd × s / m^2^ flash stimuli, where both, native cone-opsins and ReaChR are expected to be activated.

### Transmittance of native and optogenetic signals to the brain

After studying the mode of interference between natural and optogenetic responses on a retinal level, we asked if and how these events would shape cortical VEP responses. Consistent with our recordings from dark-adapted retinas we observed no difference in VEP N1 amplitudes or timing evoked by a 1 cd×s/m^2^ flash on a dark background (**Supplementary Figure 3**). We next assessed VEP responses under light adapted conditions using a flash energy of 100 cd×s/m^2^ that had evoked the optogenetic a_o_-wave on ERG. Expectably, light-adapted VEP responses were small, and it was not possible to unequivocally identify a signal in each mouse (**Figure 7 A**). Overall, P1 and N1 components of the VEP could be successfully annotated in three out of seven (42.9 %) wt mice and 9 out of 10 wt+ReaChR mice (90 %). Thus, there were too few data points in the wt cohort for and inter-group comparison. We therefore restricted ourselves to analyses within the wt+ReaChR mice only. Interestingly, as for the ERG data, an early negative deflection, sometimes just above noise level and preceding the N1-wave, could be observed in most (8/9) recordings (**Figure 7 A, red arrow**): Median time to N1 peak was 53.5 [IQR: 51.5 – 59.0] ms while N1_o_ peak occurred at 27.6 [IQR: 25.1 – 29.9] ms (n=8, **Figure 7 B**). We hypothesized that this early deflection might be the cortical analogue of the retinal a_o_-wave, i.e. a response evoked by optogenetic activation reaching the cortex earlier than the native signal. Under this hypothesis, we called the early deflection N1_o_. If this hypothesis was true, and assuming that the native and the optogenetic signal would need approximately the same time to travel to the cortex, the latency between a_o_- and N1_o_-waves should match that between a- and N1-waves on an individual level. Indeed, a_o_-N1_o_-latencies closely correlated with a-N1-latencies (R = 0.73, p = 0.038, **Figure 7C**), thus strongly indicating that N1_o_ is of optogenetic origin signifying asynchronous arrival of the native and optogenetically captured visual information at the level of the cortex.

**Figure 7:**
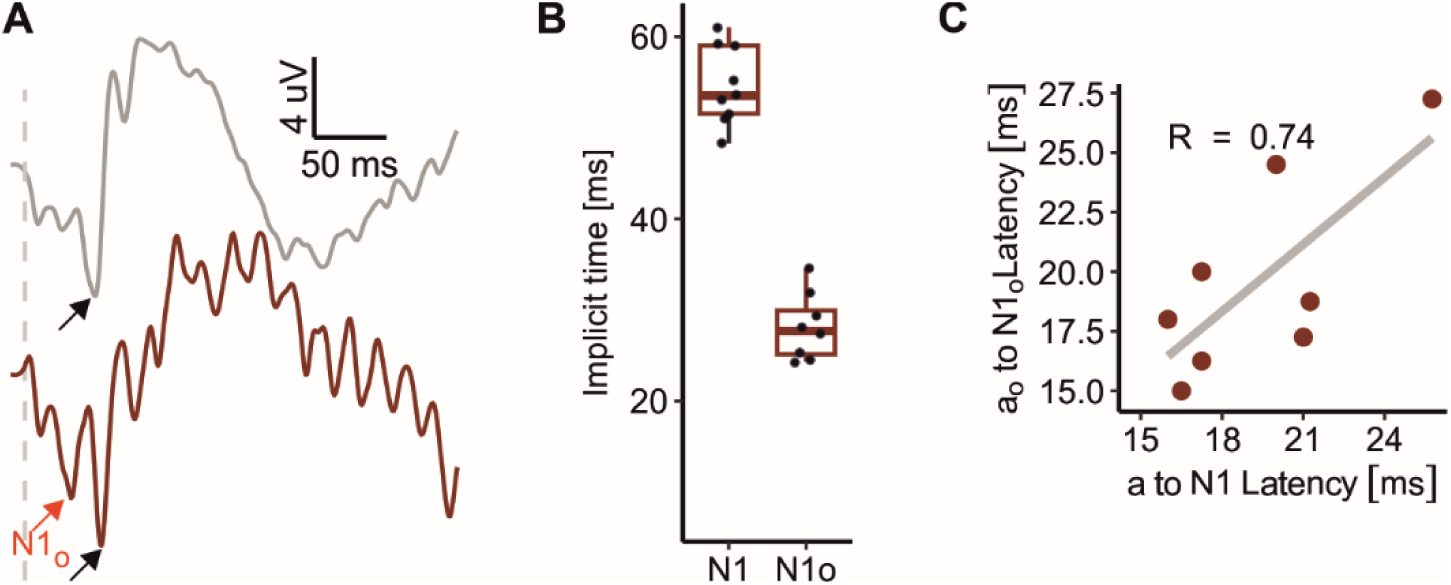
Light adapted VEP recordings. **(A)** Representative recordings from wild-type mice (grey) and ReaChR-expressing, non-degenerate littermates (dark red) obtained in response to a 100 cd × s / m^2^ flash stimulus on 30 cd / m^2^ background. Light adapted VEP amplitudes were low and signal-to-noise ratios sufficient for further analysis could only be obtained 9 out of 10 recordings from ReaChR-expressing, non-degenerate mice. Note an additional, early deflection (N1_o_, red arrow) could be observed in all but one recording from ReaChR-expressing, non-degenerate mice but not in those from wild-type mice not expressing ReaChR. Black arrow points on the N1 wave. **(B)** Summary statistics for the implicit times of ERG a and a_o_ as well as VEP N1 and N1_o_ waves. Median time to N1 peak was 27.6 [IQR: 25.1 – 29.9] ms while N1_o_ peak occurred at 27.6 [IQR: 25.1 – 29.9] ms. **(C)** Scatter plot showing the latencies from a to N1 waves versus the latencies from a_o_ to N1_o_ waves. Latencies significantly correlate indicating that the N1_o_ likely reflects the cortical response to ReaChR activation.

## DISCUSSION

In the present work, we use a model of OBC targeted optogenetic therapy to demonstrate that native (rod and cone driven) and optogenetic responses adversely interact. Optogenetic responses are enhanced in the presence of concomitant native photoreception while native responses become reduced. Moreover, optogenetic responses reach the cortex in advance of the native response and this temporal desynchrony may impact on visual perception.

The observations made herein need to be considered whenever the am is develop or test an optogenetic vision restorative therapy in individuals with residual native vision, be it to offer restorative treatment for patients with late-stage atrophic age-related macular degeneration or to pre-empt potential retinal rewiring that might hinder optimal outcomes in vision-restorative approaches.

### The a_o_ wave

Hallmark of the ERG in retinas expressing the optogenetic tool ReaChR is the a_o_-wave, a fast and early negative deflection in response to bright (100 cd×s/m^2^ ≈ 1.17 × 10^15^ photons/cm^2^/s at pupil level) flashes that we exclusively observe in mice expressing ReaChR. This deflection peaked at 8.0 [IQR: 7.1 - 8.8] ms in wt+ReaChR mice, which was substantially (and significantly) earlier than the time-to-peak we observed for wt mouse ERG a-waves. From our previous *ex-vivo* electrophysiology studies [11, 23] as well as the stimulus-response curves published for ReaChR in cell lines [20] we can expect to see ReaChR driven responses from a photon flux of 10^15^ photons/cm^2^/s and above. This fits very well to our observation that a_o_ is apparent from 100 cd×s/m^2^ onwards but is absent at lower light levels.

With the experiments performed herein, we cannot unequivocally conclude on the origins of the a_o_-wave. In theory it may be either a direct and exclusive reflection of ReaChR activation, it may carry contributions of another OBC conductance activated secondary to ReaChR-activation, or it may even have contributions from other cell types, either via synaptic mechanisms or reflex Muller-cell mediated K+ clearance. Indeed, the a_o_ implicit time of 8.0 [IQR: 7.1 - 8.8] ms seems too short to allow any chemically-synaptic transmission, ruling out the latter option. Also, we observed the onset of the a_o_-wave to be faster than could be resolved by our ERG recordings at the employed sampling rate (2 kHz) strongly suggesting that ReaChR activation is directly responsible for the upstroke of the a_o_-wave.

Activated Channelrodopsin-2-derivatives like ReaChR carry an unselective cation current and thus show a reversal potential close to 0 mV [24]. Consequently, light mediated activation of ReaChR should result in a depolarization of OBCs. Under the assumption that the native b-wave would be directly carried by OBC depolarization, b- and a_o_-wave should have the same polarity. It might therefore seem unintuitive that the ERG correlate of optogenetic activation we observe is a negative wave. Though we did not dissect the cellular basis of this response polarity in detail, our observation is in line with several others previously reported ERG responses in retina-degenerate optogenetically treated animals [10, 25]. Indeed, there is an ongoing debate that reflective Muller-cell depolarization may have a substantial contribution to the b-wave [26]. This does not only offer a potential explanation for the inverse polarity of b- and a_o_-wave, it could moreover explain why in rd1+ReaChR we do see a slow peak following the a_o_-wave that approximately resembles the kinetics of the native b-wave. Thus, the polarity of the optogenetic a_o_-wave is not only in line with previous reports, it also fits into the current model of the origins the ERG components in the mammalian retina.

### Larger optogenetic responses in the non-degenerate retina

Comparing our data from retina-degenerate rd1+ReaChR and non-degenerate wt+ReaChR mice we find that optogenetic responses (as measured by a_o_-wave amplitude) are substantially larger in non-degenerate retinas. It might seem an intuitive explanation that in absence of glutamatergic input in the degenerate retina OBCs rest at more depolarized potentials, therefore field potential changes caused by ReaChR activation would be smaller. Yet, it has been frequently observed that in absence of glutamatergic input OBCs indeed hyperpolarize [27–29], possibly due to Trpm1 down-regulation [27, but see also: 30]. Thus, it is more likely that other factors, e.g. differences in OBC membrane resistance, underly the attenuated response amplitudes. In agreement with the concept that “unstimulated” ReaChR might attenuate OBC membrane resistance are also the observed tendency towards longer b-wave implicit times in ReaChR mice. Future studies likely including single-unit electrophysiological studies will be required to further elucidate the mechanisms underlying our observation and might pave the way towards engineering optogenetic tool that overcome this effect.

### Dampening of the cone-mediated b-wave

Another observation made in this study is the dampening of the b-wave under lighting conditions eliciting cone responses that are below the apparent “ReaChR activation threshold”. For interpretation it needs to be kept in mind that ReaChR is an ion channel that briefly opens following the absorption of a photon. Thus, there is no “activation threshold” per se but rather a threshold from which the number of ReaChR ion channel openings and size of ReaChR-currents are sufficient to change the signalling behaviour of a cell. Consequently, even light intensities too low to evoke an a_o_-wave may already affect the electrical properties of OBCs and we speculate that this is the most likely cause of the observed b-wave dampening seen in non-degenerate ReaChR expressing wt+ReaChR mice.

### Native and optogenetic responses at the cortex

Visual perception is created at cortical level. To assess how the interference of native and optogenetic retina responses observed by ERG would affect the signal arriving at the visual cortex we additionally performed flash VEP recordings. While we only observed a single negative deflection in response to a 100 cd×s/m^2^ flash in mice not expressing ReaChR, in wt+ReaChR mice we observed an additional negative deflection prior to the native N1 wave. This suggests that optogenetic visual responses arrive at cortical level slightly prior to the native visual responses. This could be mechanistically explained as the cone-OBC-borne optogenetic responses have to pass one synapse less than the native responses [31] and the fast kinetics of the light gated ReaChR channel are not dependent on a signalling cascade. For a partially sighted individual receiving optogenetic gene therapy, this asynchrony in signal arrival could result in a temporally scrambled perception. It is worth noting that, with the subdermal electrode placement used in this study, we were technically near the detection limit for these dual-dip VEP responses. Thus, further studies will be required to dissect mixed native and optogenetic visual responses on cortical level in more detail. Indeed, it is conceivable that the brain would be able to adapt to such asynchronous signals and still make sense of this modified sensory input. Nevertheless, these observations need to be taken into consideration when planning clinical trials in partially sighted patients, and further preclinical studies will be needed to optimize optogenetic therapies in this regard.

### Segregation of optogenetic and native responses

Identification and segregation of optogenetic and native visual responses in this study was mainly achieved by comparing between different genotypes. We were also able to differentially stimulate native and optogenetic components by using stimulus paradigms that took advantage of the distinct spectral sensitivities of ReaChR and the native cone opsins. As sensitivity spectra of cone opsins and ReaChR overlap, a pure activation of native vs. optogenetic photoreceptors has not been possible. Still, we could use this stimulus paradigm to show that optogenetic activity per se and not just temporal overlap with native responses is responsible for the damping of b-wave amplitudes. Technical refinement of our experimental procedure, e.g. using silent substitution or *in-vivo* pharmacology may enable an isolated assessment of native and optogenetic responses in the same animal in future.

### Specific relevance for treating macular disease and retinitis pigmentosa

The main focus of optogenetic visual restorative approaches thus far has been on restoring vision in terminally blind patients suffering from retinitis pigmentosa [8]. While this is certainly the patient collective that has most to benefit from successful optogenetic vision restoration, the number of patients suffering from macular degenerations, in particular age-related macular degeneration, far outnumbers those suffering from retinitis pigmentosa [3, 4]. It is therefore important to understand how optogenetic vision restoration could function in macular diseases, where areas of atrophic and intact retina coexist in close proximity (**Figure 1**) [32, 33]. In this regard, the results presented herein indicate that efforts should be made to develop approaches that restrict the expression of the optogenetic tool to the area affected by retinal atrophy in order to avoid interference with native residual vision. Locally restricted optogenetic activation could already be achieved using signal pre-processing goggles. Yet, we also observe that the presence of the optogenetic tool ReaChR in OBCs dampens native ERG responses to moderately bright ambient light level stimuli. In functional terms, this might mean that the residual native vision in a macular degeneration patient would be hindered by an optogenetic gene therapy. Obviously, this would hopefully be outweighed by the functional benefit these patients would experience by the optogenetic restoration of vision in the atrophic/degenerate areas. Yet, as the functional benefit is subject to research, this consideration needs to be included when planning clinical trials. Notably, a first clinical trial aiming on optogenetic vision restoration in patients suffering from Stargardt’s macular dystrophy is ongoing (NCT05417126, [34]). Trial reports in scientific outlets in future may serve to better estimate the functional gain or possibly the payload of optogenetic interference that can be expected in this group of patients.

Conversely, several clinical trials aiming to establish an optogenetic gene therapy for patients blinded from retinitis pigmentosa have already been (and are currently being) conducted. While providing valuable insights [9], they have not yet been able to achieve the desired gain in functional vision [Reviewed in: 8]. Among several possible explanations, one is that trails have been conducted in terminally blind (“no light perception”) individuals with a history of inner retinal remodelling spanning years or even decades, which may hinder optimal functional outcomes [Reviewed in: 8]. Consequently, optogenetic treatment may be more effective if applied earlier in the course of disease, before significant remodelling has taken place and thus probably also at a time when the patient still has some level of remaining native vision. Considerations to be made are analogous to those described for macular diseases above. Yet an additional level of complexity is added by the fact in retinitis pigmentosa cone vision is preserved longest, and we have observed that it is cone-, rather than rod-vision, that gets functionally affected by optogenetic vision.

### Limitations

In this study we have used a transgenic mouse line expressing our optogenetic tool in (virtually all) OBCs. Obviously, this deviates from a clinically translatable Adeno-associated virus mediated optogenetic gene therapy, where expectably the transfection rates are clearly below 100% and the cell type tropism might be less uniform than in our transgenic approach. Thus, the exact pattern of interaction as well as the degree might differ from what we observed here. Yet, complete, uniform and cell type-specific transgene delivery would be required to achieve optimal performance in terms of vision restoration in the blind retina and consequently be the aim of any virally delivered optogenetic gene therapy.

Recording of mouse VEPs under light-adapted conditions using subdermal electrodes is generally a challenge, as these are low in amplitude and we have been facing similar challenges resulting in only moderate signal-to-noise ratios for these recordings. This should be taken into consideration when interpreting the VEP data presented herein. Future studies employing transcranial VEP or alternative, spatially resolved, approaches like calcium imaging or functional ultrasound [35], may provide deeper insights into the cortical-level patterns of interference.

### Conclusion

Overall, in the present work we performed ERG and VEP recordings to dissect how optogenetic and native vision interact. Our key findings are that optogenetic responses are larger in healthy retinas retaining rods and cones, that native cone-driven responses are dampened by optogenetic treatment and that native and optogenetic responses arrive asynchronously at the cortex. These findings should be taken into consideration when planning future clinical trials for patients with residual visual function and may direct future research to optimize optogenetic approaches for visual restoration on preclinical level.

## MATERIAL AND METHODS

### Animal studies

All procedures were performed with the approval of the Giessen Regional Council Animal Health Authority (File No: G93/2022) and in accordance with the ARVO Statement for the Use of Animals in Ophthalmic and Vision Research. Mice were housed under a twelve-hour light / twelve-hour dark cycle with food and water available *ad libitum*.

This study uses the ReaChR Grm6 rd strain, previously described in Rodgers et al. [11], created by breeding Grm6^iCre/WT^ (MGI:4411993, kindly shared by Robert Duvoisin, Oregon Health and Science University, USA) with ReaChR-mCitrine mice (MGI: 5605725) obtained from Jax laboratories (#026294). Mice of this strain were kept heterozygous for Grm6 iCre and homozygous for Pde6b and ReaChR. Only mice heterozygous for Grm6 iCre were used in this study (“Rd1+ReaChR”). We also created a new strain that was homozygous for the wild-type Pde6b allele by back-crossing against C57BL/6J purchased from Charles River (Sulzfeld, Germany). Mice from this strain used in this study were either heterozygous for Grm6 iCre (“wt+ReaChR”) or homozygous wild-type at the Grm6 iCre locus (“wt”). Genotyping was performed as described elsewhere [11] but using DreamTaq PCR Master Mix (Thermo Fisher, Waltham, MA, USA).

### Electrophysiological Recordings

Electroretinogram recordings were performed at the age of 9-12 weeks usually at Zeitgeber time 3-6h under general anaesthesia. General anaesthesia was induced and maintained with isoflurane (Baxter, Deerfield, IL, USA), using approximately 1% isoflurane in 0.2–0.5 L/min O₂ for maintenance. The anaesthesia was delivered via a Univentor 410-Q vaporizer (UNO Roestvaststaal BV, Zevenaar, The Netherlands). Pupil dilation was achieved through the local application of Tropicamide (1%) and Phenylephrine (2.5%) eye drops. Electroretinograms were recorded using a Celeris Rodent ERG system (Diagnosys LLC, Lowell, MA, USA) employing an integrated light guide stimulator and electrode. Oxybuprocaine hydrochloride (4 mg/ml) was installed to the eye before placement of the stimulator/electrodes. Data acquisition was performed using Espion software (Diagnosys) and digitized at a sampling rate of 2000 Hz. Unless stated otherwise, stimulus protocols used were designed reflecting ISCEV standards for ERG and VEP, respectively [36, 37]. Mice were dark-adapted for at least 20 min before the beginning of the recording and thereafter only handled under dim red light. Before the beginning of the light adapted recordings, mice were exposed to 30 cd/m^2^ for at least 8 minutes. All light-adapted stimuli were then delivered onto the same 30 cd/m^2^ background. Flash stimulus duration was 4 ms maximum. Flash ERG recordings were band-pass filtered at 0.125 – 300 Hz. and VEP recordings were band-pass filtered at 3 – 100 Hz, with the lower edge of the filter slightly deviating from ISCEV standards as this resulted in a clearer demarcation of the ReaChR VEP responses. Recordings were exported from the Espion database into CSV files using the software’s inbuilt function as described previously [38] and then analysed using the ERGtools2 package [38] (https://github.com/moritzlindner/ERGtools2) for R [39]. For ERG recordings a-wave amplitudes are reported as absolute values, while b wave amplitudes are reported relative to the a-wave. Similarly, for VEP recordings N1 wave amplitudes are presented relative to the preceding P1 wave.

### Immunohistochemistry

For histological analysis, tissue was collected and processed as described earlier in detail [40]. In brief, mice were culled by decapitation under isoflurane anaesthesia and eyes were enucleated. Eyes were fixed in 4% methanol-free paraformaldehyde (Thermo Fisher) in phosphate buffered saline (PBS) and embedded into Optimal Cutting Temperature (OCT) medium (VWR, Lutterworth, UK). 18 µm tissue sections were prepared using a CM1850 Cryotome (Leica, Wetzlar, Germany). Retinal cryosections sections were blocked in PBSTX-0.2 with 10% normal donkey or goat serum (Sigma Aldrich, St. Louis, USA). Sections were then incubated with primary antibodies for 24 h at 4 °C and with secondary antibodies for 2 hours at room temperature. All antibodies were diluted in PBSTX-0.2 containing 2.5% normal donkey serum. The following primary antibodies were used: Chicken anti-GFP (also recognizing mCitrine; AVES Labs, Tigard, OR, USA, GFP-1020; RRID: AB_2307313; Dilution: 1:500), rabbit anti-Cone arrestin (Merck, Darmstadt, Germany, AB15282; RRID: AB_1163387; Dilution: 1:500). PNA Lectin conjucated to Alexa 647 was purchased from Thermo Fisher (Dilution 1:50).

Image acquisition and analysis was performed using an upright LSM 710 laser scanning confocal microscope (Carl Zeiss Meditec, Oberkochen, Germany) for acquisition and ImageJ [41] software for analysis as described before [42].

### Statistical Analysis and Visualization

Statistical analysis and data visualization was performed using R[39] core packages, ERGtools2 [38] and ggplot2 [43]. Boxplots shown follow the definition of Tukey [44], with the thick line representing the median and the upper and lower hinges 1^st^ and 3^rd^ quartile, respectively. Line diagrams represent group means and error bars represent the standard error of the mean. Unpaired t-tests or Wilcoxon tests were used for inferential statistics after testing for normality using Shapiro-Wilk test. Functions used for conversion between light units are available at https://github.com/moritzlindner/lindnerlab/.

## ACKNOWLEDGEMENTS

We thank Prof. Burkhard Schütz and the Dept. of Anatomy (University of Marburg) for providing access to their cryotome, Robert Duvoisin (Oregon Health & Science University) for kindly sharing the Grm6Cre mouse line and Mrs. Irina Bogun for supreme technical assistance.

## DECLARATIONS

### Author Contributions

Participated in research design: ML, MJG

Conducted experiments: ML, EC, SH, NX, SB

Performed data analysis: ML

Wrote or contributed to the writing of the manuscript: ML, SH, JR, MJG, MWH

### Data Availability

The datasets generated during and/or analysed during the current study are available from the corresponding author on reasonable request.

### Funding

Supported by the German Research Foundation (LI 2846/5-1 and LI 2846/6-1 to ML) and the ProRetina Foundation (Pro-Re/Projekt/Gi-Wh-Li.04-2021 to ML and MJG).

### Declaration of Interest

ML has received Grants from Bayer Healthcare outside the submitted work.

### Ethics approval

This study did not involve human participants. Animal work was performed with approval of the relevant authorities and in accordance with the institutional Ethics Guidelines of Animal Care. Further details are provided in the Methods section.

### Consent to publish

Not applicable.

## SUPPLEMENTARY TABLES

**Supplementary Table 1:**
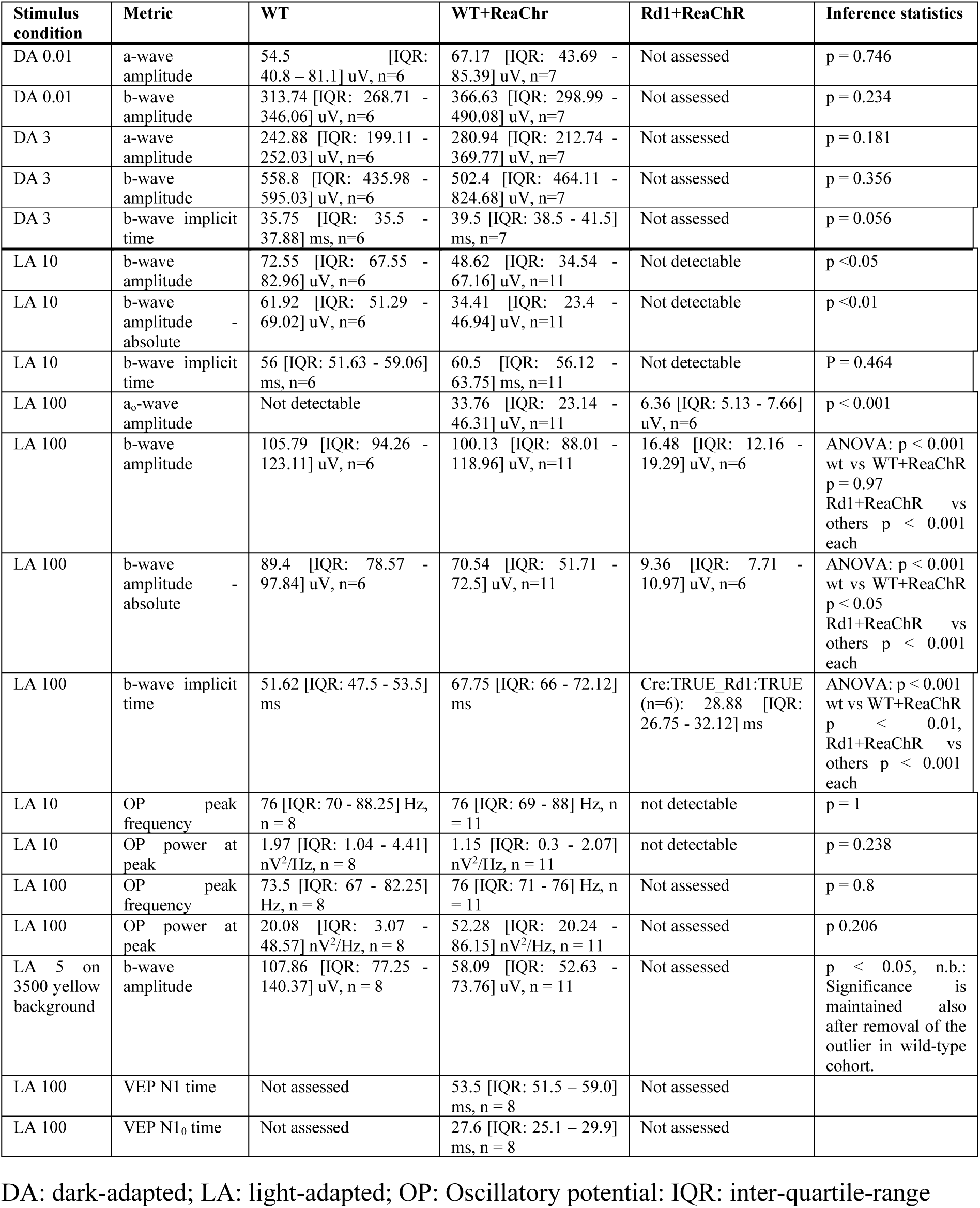
Summary of metrics presented in this paper.

## SUPPLEMENTARY FIGURES

**Supplementary Figure 1:**
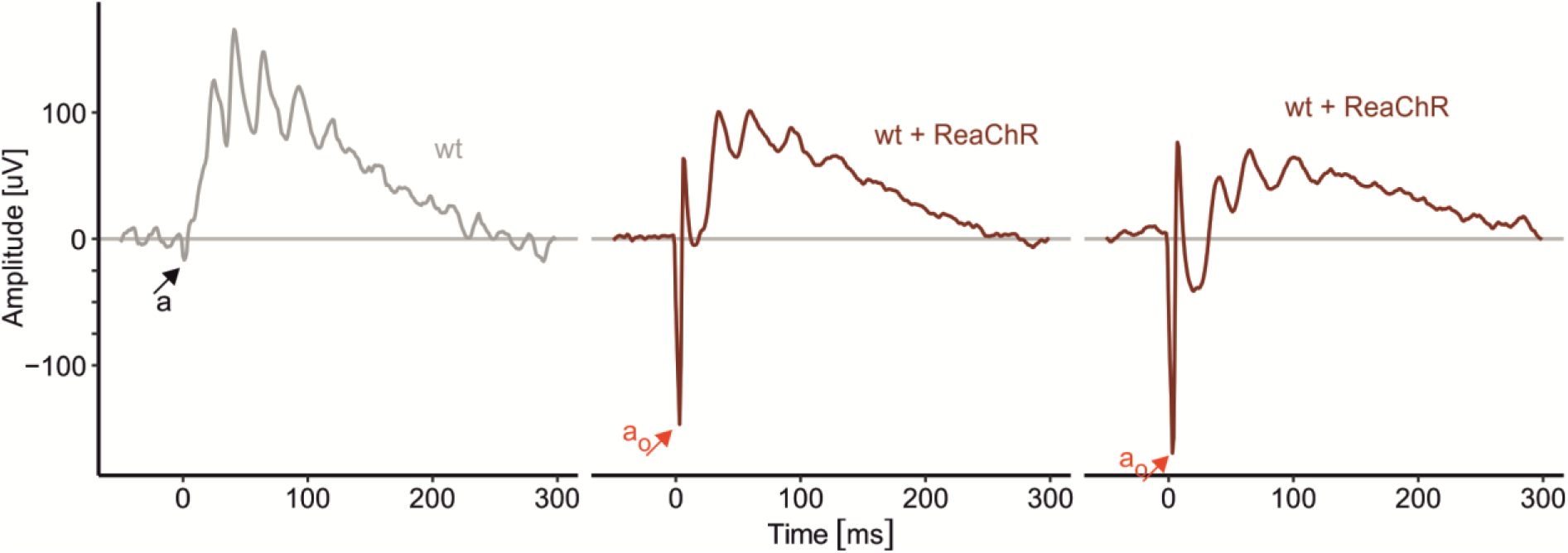
Exemplary recordings from three mice (one wild-type, left, and two ReaChR expressing, non-degenerate) obtained under light-adapted conditions in response to a 900 cd × s / m^2^ flash stimulus. Early (Implicit time: ∼7 ms) a_o_ waves could only be observed in ReaChR expressing, non-degenerate mice.

**Supplementary Figure 2:**
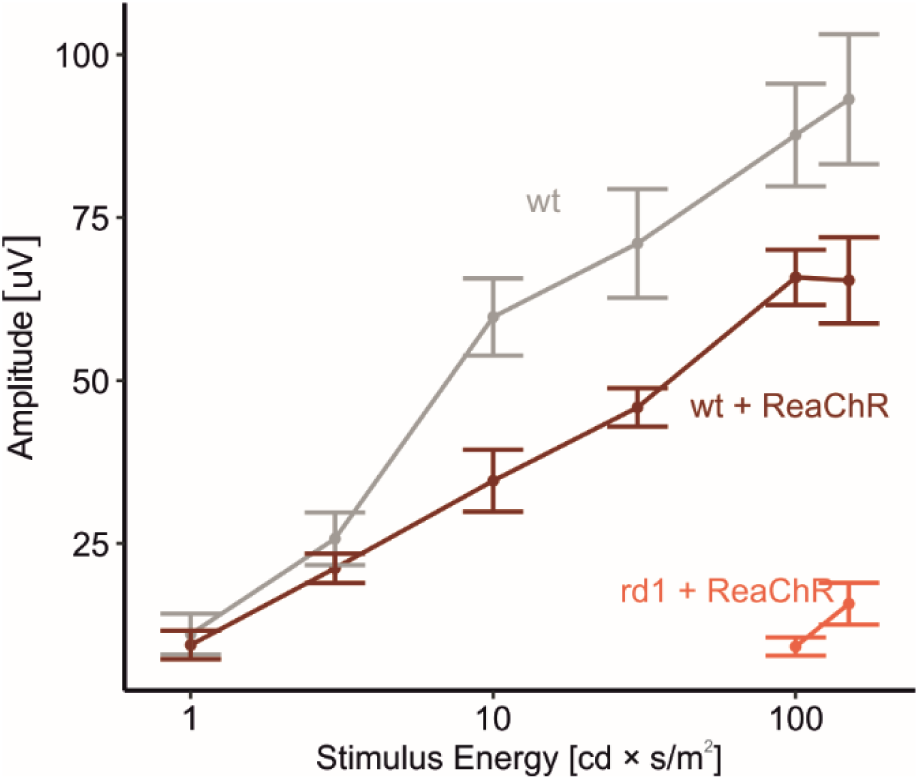
Stimulus – Response curves for absolute b-wave amplitudes. Same data as shown in Figure 4 E, but amplitudes measured as absolute values instead of relative to the a-wave.

**Supplementary Figure 3:**
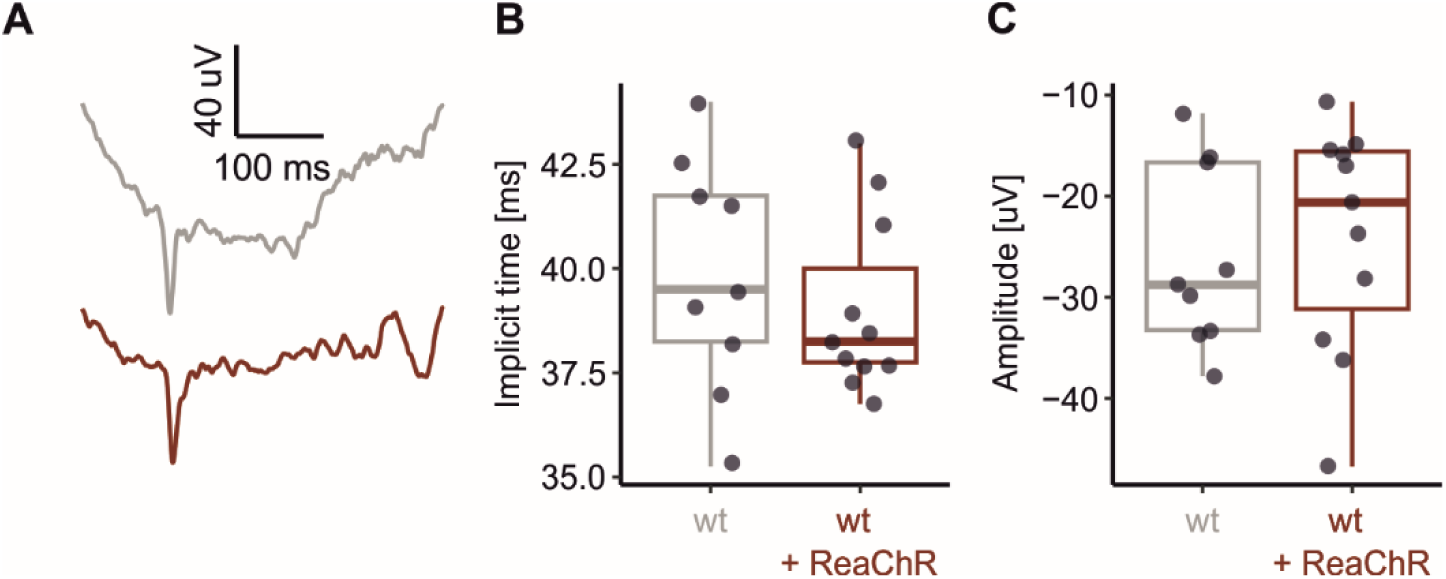
Dark-adapted (rod-dominated) VEP recordings are not altered in mice expressing ReaChR in OBC. **(A)** Representative recordings obtained in response to a 1 cd × s / m^2^ flash stimulus from wild-type mice (grey) and ReaChR-expressing, non-degenerate littermates (dark red). **(B, C)** Summary statistics for N1-wave implicit times (B) and amplitudes (C), respectively.

## REFERENCES

1. Galvin, O., Chi, G., Brady, L., Hippert, C., Del Valle Rubido, M., Daly, A., and Michaelides, M. (2020). The Impact of Inherited Retinal Diseases in the Republic of Ireland (ROI) and the United Kingdom (UK) from a Cost-of-Illness Perspective. Clin Ophthalmol 14, 707–719.

2. Chuvarayan, Y., Finger, R.P., and Koberlein-Neu, J. (2020). Economic burden of blindness and visual impairment in Germany from a societal perspective: a cost-of-illness study. Eur J Health Econ 21, 115–127.

3. Cross, N., van Steen, C., Zegaoui, Y., Satherley, A., and Angelillo, L. (2022). Retinitis Pigmentosa: Burden of Disease and Current Unmet Needs. Clin Ophthalmol 16, 1993–2010.

4. World Health Organization (2019). World Report on Vision, (Geneva).

5. Sullivan, L.S.D., S.P. (2024). retnet.org.

6. Garg, A., Nanji, K., Tai, F., Phillips, M., Zeraatkar, D., Garg, S.J., Sadda, S.R., Kaiser, P.K., Guymer, R.H., Sivaprasad, S., et al. (2024). The effect of complement C3 or C5 inhibition on geographic atrophy secondary to age-related macular degeneration: A living systematic review and meta-analysis. Surv Ophthalmol 69, 349–361.

7. Lorenz, B., Kunzel, S.H., Preising, M.N., Scholz, J.P., Chang, P., Holz, F.G., and Herrmann, P. (2024). Single Center Experience with Voretigene Neparvovec Gene Augmentation Therapy in RPE65 Mutation-Associated Inherited Retinal Degeneration in a Clinical Setting. Ophthalmology 131, 161–178.

8. Lindner, M., Gilhooley, M.J., Hughes, S., and Hankins, M.W. (2022). Optogenetics for visual restoration: From proof of principle to translational challenges. Prog Retin Eye Res 91, 101089.

9. Sahel, J.A., Boulanger-Scemama, E., Pagot, C., Arleo, A., Galluppi, F., Martel, J.N., Esposti, S.D., Delaux, A., de Saint Aubert, J.B., de Montleau, C., et al. (2021). Partial recovery of visual function in a blind patient after optogenetic therapy. Nat Med 27, 1223–1229.

10. Kralik, J., and Kleinlogel, S. (2021). Functional Availability of ON-Bipolar Cells in the Degenerated Retina: Timing and Longevity of an Optogenetic Gene Therapy. Int J Mol Sci 22.

11. Rodgers, J., Hughes, S., Lindner, M., Allen, A.E., Ebrahimi, A.S., Storchi, R., Peirson, S.N., Lucas, R.J., and Hankins, M.W. (2023). Functional integrity of visual coding following advanced photoreceptor degeneration. Curr Biol 33, 474–486 e475.

12. Busskamp, V., Duebel, J., Balya, D., Fradot, M., Viney, T.J., Siegert, S., Groner, A.C., Cabuy, E., Forster, V., Seeliger, M., et al. (2010). Genetic reactivation of cone photoreceptors restores visual responses in retinitis pigmentosa. Science 329, 413–417.

13. Mace, E., Caplette, R., Marre, O., Sengupta, A., Chaffiol, A., Barbe, P., Desrosiers, M., Bamberg, E., Sahel, J.A., Picaud, S., et al. (2015). Targeting channelrhodopsin-2 to ON-bipolar cells with vitreally administered AAV Restores ON and OFF visual responses in blind mice. Mol Ther 23, 7–16.

14. Cehajic-Kapetanovic, J., Eleftheriou, C., Allen, A.E., Milosavljevic, N., Pienaar, A., Bedford, R., Davis, K.E., Bishop, P.N., and Lucas, R.J. (2015). Restoration of Vision with Ectopic Expression of Human Rod Opsin. Curr Biol 25, 2111–2122.

15. Bi, A., Cui, J., Ma, Y.P., Olshevskaya, E., Pu, M., Dizhoor, A.M., and Pan, Z.H. (2006). Ectopic expression of a microbial-type rhodopsin restores visual responses in mice with photoreceptor degeneration. Neuron 50, 23–33.

16. Berry, M.H., Holt, A., Salari, A., Veit, J., Visel, M., Levitz, J., Aghi, K., Gaub, B.M., Sivyer, B., Flannery, J.G., et al. (2019). Restoration of high-sensitivity and adapting vision with a cone opsin. Nat Commun 10, 1221.

17. Pfeiffer, R.L., Marc, R.E., and Jones, B.W. (2020). Persistent remodeling and neurodegeneration in late-stage retinal degeneration. Prog Retin Eye Res 74, 100771.

18. Lindner, M., Kosanetzky, S., Pfau, M., Nadal, J., Gordt, L.A., Schmitz-Valckenberg, S., Schmid, M., Holz, F.G., Fleckenstein, M., and Group, F.A.-S. (2018). Local Progression Kinetics of Geographic Atrophy in Age-Related Macular Degeneration Are Associated With Atrophy Border Morphology. Invest Ophthalmol Vis Sci 59, AMD12-AMD18.

19. Lindner, M., Lambertus, S., Mauschitz, M.M., Bax, N.M., Kersten, E., Luning, A., Nadal, J., Schmitz-Valckenberg, S., Schmid, M., Holz, F.G., et al. (2017). Differential Disease Progression in Atrophic Age-Related Macular Degeneration and Late-Onset Stargardt Disease. Invest Ophthalmol Vis Sci 58, 1001–1007.

20. Lin, J.Y., Knutsen, P.M., Muller, A., Kleinfeld, D., and Tsien, R.Y. (2013). ReaChR: a red-shifted variant of channelrhodopsin enables deep transcranial optogenetic excitation. Nat Neurosci 16, 1499–1508.

21. Morgans, C.W., Zhang, J., Jeffrey, B.G., Nelson, S.M., Burke, N.S., Duvoisin, R.M., and Brown, R.L. (2009). TRPM1 is required for the depolarizing light response in retinal ON-bipolar cells. Proc Natl Acad Sci U S A 106, 19174–19178.

22. Wachtmeister, L. (1998). Oscillatory potentials in the retina: what do they reveal. Prog Retin Eye Res 17, 485–521.

23. Lindner, M., Gilhooley, M.J., Peirson, S.N., Hughes, S., and Hankins, M.W. (2021). The functional characteristics of optogenetic gene therapy for vision restoration. Cell Mol Life Sci 78, 1597–1613.

24. Feldbauer, K., Zimmermann, D., Pintschovius, V., Spitz, J., Bamann, C., and Bamberg, E. (2009). Channelrhodopsin-2 is a leaky proton pump. Proc Natl Acad Sci U S A 106, 12317–12322.

25. Nikonov, S., Aravand, P., Lyubarsky, A., Nikonov, R., Luo, A.J., Wei, Z., Maguire, A.M., Phelps, N.T., Shpylchak, I., Willett, K., et al. (2022). Restoration of Vision and Retinal Responses After Adeno-Associated Virus-Mediated Optogenetic Therapy in Blind Dogs. Transl Vis Sci Technol 11, 24.

26. Bhatt, Y., Hunt, D.M., and Carvalho, L.S. (2023). The origins of the full-field flash electroretinogram b-wave. Front Mol Neurosci 16, 1153934.

27. Xu, Y., Dhingra, A., Fina, M.E., Koike, C., Furukawa, T., and Vardi, N. (2012). mGluR6 deletion renders the TRPM1 channel in retina inactive. J Neurophysiol 107, 948–957.

28. Borowska, J., Trenholm, S., and Awatramani, G.B. (2011). An intrinsic neural oscillator in the degenerating mouse retina. J Neurosci 31, 5000–5012.

29. Gregg, R.G., Kamermans, M., Klooster, J., Lukasiewicz, P.D., Peachey, N.S., Vessey, K.A., and McCall, M.A. (2007). Nyctalopin expression in retinal bipolar cells restores visual function in a mouse model of complete X-linked congenital stationary night blindness. J Neurophysiol 98, 3023–3033.

30. Gilhooley, M.J., Hickey, D., Lindner, M., Palumaa, T., Hughes, S., Peirson, S.N., MacLaren, R.E., and Hankins, M.W. (2021). ON-bipolar cell gene expression during retinal degeneration: Implications for optogenetic visual restoration. Exp Eye Res, 108553.

31. Wassle, H. (2004). Parallel processing in the mammalian retina. Nat Rev Neurosci 5, 747–757.

32. Pfau, M., Muller, P.L., von der Emde, L., Lindner, M., Moller, P.T., Fleckenstein, M., Holz, F.G., and Schmitz-Valckenberg, S. (2020). Mesopic and Dark-Adapted Two-Color Fundus-Controlled Perimetry in Geographic Atrophy Secondary to Age-Related Macular Degeneration. Retina 40, 169–180.

33. Lindner, M., Pfau, M., Czauderna, J., Goerdt, L., Schmitz-Valckenberg, S., Holz, F.G., and Fleckenstein, M. (2019). Determinants of Reading Performance in Eyes with Foveal-Sparing Geographic Atrophy. Ophthalmol Retina 3, 201–210.

34. Mahajan, V.B. (2024). Longitudinal BCVA Analysis of Patients With Stargardt Disease and Macular Degeneration Treated With MCO-010, a Mutation-Agnostic Optogenetic Therapy: 48-Week Results From a Phase 2a Clinical Trial (STARLIGHT). Investigative Ophthalmology & Visual Science 65, 5266–5266.

35. Edelman, B.J., Siegenthaler, D., Wanken, P., Jenkins, B., Schmid, B., Ressle, A., Gogolla, N., Frank, T., and Mace, E. (2024). The COMBO window: A chronic cranial implant for multiscale circuit interrogation in mice. PLoS Biol 22, e3002664.

36. Odom, J.V., Bach, M., Brigell, M., Holder, G.E., McCulloch, D.L., Mizota, A., Tormene, A.P., and International Society for Clinical Electrophysiology of, V. (2016). ISCEV standard for clinical visual evoked potentials: (2016 update). Doc Ophthalmol 133, 1–9.

37. McCulloch, D.L., Marmor, M.F., Brigell, M.G., Hamilton, R., Holder, G.E., Tzekov, R., and Bach, M. (2015). ISCEV Standard for full-field clinical electroretinography (2015 update). Doc Ophthalmol 130, 1–12.

38. Lindner, M. (2024). The ERGtools2 package: A Toolset for Processing and Analysing Visual Electrophysiology Data. bioRxiv, 2024.2008.2027.609856.

39. R Core Team (2018). R: A Language and Environment for Statistical Computing. (Vienna, Austria: R Foundation for Statistical Computing).

40. Kinder, L., and Lindner, M. (2024). Expression of Osteopontin in M2 and M4 intrinsically photosensitive retinal ganglion cells. bioRxiv, 2024.2010.2031.621275.

41. Schindelin, J., Arganda-Carreras, I., Frise, E., Kaynig, V., Longair, M., Pietzsch, T., Preibisch, S., Rueden, C., Saalfeld, S., Schmid, B., et al. (2012). Fiji: an open-source platform for biological-image analysis. Nat Methods 9, 676-682.

42. Lindner, M., Gilhooley, M.J., Palumaa, T., Morton, A.J., Hughes, S., and Hankins, M.W. (2020). Expression and Localization of Kcne2 in the Vertebrate Retina. Invest Ophthalmol Vis Sci 61, 33.

43. Wickham, H. (2016). ggplot2: Elegant Graphics for Data Analysis. (Springer-Verlag New York).

44. Robert Mcgill, J.W.T., and Wayne, A.L. (1978). Variations of Box Plots. The American Statistician 32, 12--16.

